# Growth Hormone Excess Drives Liver Aging via increased Glycation stress

**DOI:** 10.1101/2025.01.06.631635

**Authors:** Parminder Singh, Anil Gautam, Marissa Trujillo, James Galligan, Lisa Hensley, Pankaj Kapahi, Andrzej Bartke

## Abstract

Growth hormone (GH) plays a crucial role in various physiological functions, with its secretion tightly regulated by complex endocrine mechanisms. Pathological conditions such as acromegaly or pituitary tumors result in elevated circulating GH levels, which have been implicated in a spectrum of metabolic disorders, potentially by regulating liver metabolism. In this study, we focused on the liver, a key organ in metabolic regulation and a primary target of GH, to investigate the impact of high circulating GH on liver metabolism. We used bovine GH overexpressing transgenic (bGH-Tg) mice to conduct a comprehensive transcriptomic analysis of hepatic tissues. The bGH-Tg mouse livers exhibit dysregulated fatty acid metabolism and heightened inflammatory responses. Notably, the transcriptomic profile of young bGH-Tg mouse livers resembled that of aged livers and displayed markers of increased cellular senescence. Furthermore, these mice exhibited a significant accumulation of advanced glycation end products (AGEs). Intervention with glycation-lowering compounds effectively reversed the insulin resistance and aberrant transcriptomic signatures in the liver that are associated with elevated GH levels. These findings underscore the potential therapeutic value of glycation-lowering agents in mitigating the deleterious effects of chronic GH overexpression.

**Highlights:**

- Overexpression of bovine growth hormone impacts transcriptional changes in liver fat metabolism and inflammatory response in mice.
- High circulating growth hormone leads to transcriptional changes that suggest enhanced liver aging and induce cellular senescence.
- Detoxification pathways in bGH-Tg mice are inhibited, leading to the accumulation of Advanced Glycation End (AGE) products.
- Glycation-lowering compounds can mitigate pathologies associated with high GH levels.

## Introduction

Growth hormone (GH) is a pivotal endocrine regulator of growth and metabolism, orchestrating a range of physiological functions, including protein synthesis, lipolysis, and glucose homeostasis[1–3]. The secretion of GH is tightly controlled by a complex interplay of hormonal signals from the hypothalamus and pituitary gland, as well as feedback mechanisms involving insulin-like growth factor 1 (IGF-1)[4]. Dysregulation of GH secretion can lead to pathological conditions such as acromegaly and pituitary adenomas, characterized by excessively high circulating GH levels[5,6]. These conditions have been linked to an array of metabolic disturbances, including insulin resistance, dyslipidemia, and increased cardiovascular risk.

GH, which profoundly influences the liver, a central organ in metabolic homeostasis. GH directly affects hepatic function, modulating processes such as gluconeogenesis, fatty acid oxidation, and the production of IGF-1[1,7]. However, the specific molecular mechanisms by which elevated GH levels impact liver metabolism and contribute to hepatic pathologies remain incompletely understood. Previous studies have suggested that chronic GH overexpression can induce hepatic steatosis and inflammation, yet the detailed transcriptomic changes have not been fully elucidated.

To address these gaps, we utilized bovine growth hormone transgenic (bGH-Tg) mice, a model for chronic GH overexpression. By performing comprehensive transcriptomic analyses of liver tissues from these mice, we aimed to elucidate the effects of high circulating GH levels on hepatic metabolism and inflammatory responses. Our findings indicate that bGH-Tg mice exhibit significant perturbations in fatty acid metabolism and activation of inflammatory pathways. Remarkably, the liver transcriptomic profile of young bGH-Tg mice mirrors that of the aged liver, with pronounced signs of cellular senescence and AGE accumulation. The accumulation of AGEs, known to be associated with various chronic diseases, suggests a potential therapeutic target. AGEs are formed through non-enzymatic glycation of proteins and lipids, leading to altered cellular function and signaling. Given the observed accumulation of AGEs in bGH-Tg mice, we investigated the therapeutic potential of glycation-lowering compounds. Our results demonstrate that treatment with these compounds can reverse the deleterious transcriptomic changes in the liver and mitigate multiple pathologies associated with elevated GH levels.

In this study, we provide a detailed analysis of the molecular changes induced by high circulating GH in the liver. We propose glycation-lowering interventions as a promising strategy to combat the associated metabolic disorders. These findings contribute to a deeper understanding of the hepatic effects of GH and highlight novel therapeutic avenues for conditions characterized by GH dysregulation.

## Results

### Effect of GH overexpression on the liver transcriptome

Previous studies have established that chronic exposure to elevated growth hormone (GH) levels can precipitate hepatic and other pathology in bGH-Tg mice [8–10]. To elucidate the molecular underpinnings of GH-induced liver dysfunction, we conducted bulk RNA sequencing of liver tissues from bGH-Tg mice and age-matched wild-type (WT) controls. Our analysis identified significant segregation and dysregulation in gene expression, with 1193 genes downregulated and 3169 genes upregulated in bGH-Tg mice compared to WT controls (Fig. 1a-b).

**Figure 1.**
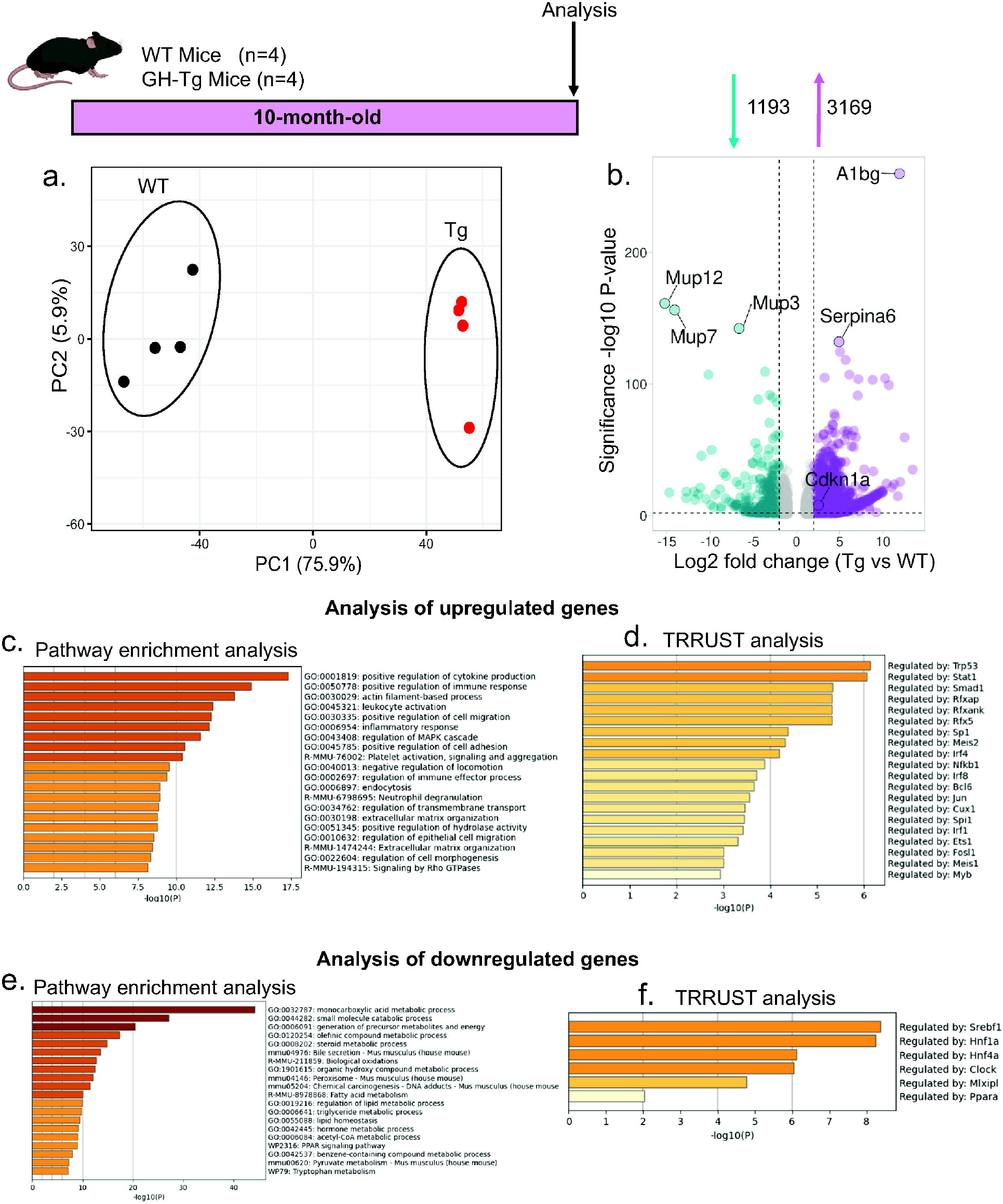
Transcriptomic Alterations in the Liver of bGH-Tg Mice. (a) Principal component analysis of RNA-seq data showing distinct clustering of bGH-Tg mice and WT controls. (b) Volcano plot highlighting differentially expressed genes (DEGs) in bGH-Tg mice compared to WT controls (adjusted p < 0.05). (c) Pathway enrichment analysis of upregulated genes reveals activation of inflammatory and immune pathways, including cytokine production and leukocyte activation. (d) TRRUST analysis identifies transcription factors such as NF-kB1 and STAT1 driving inflammatory responses in bGH-Tg mice. (e) Downregulated genes are enriched in pathways critical for lipid metabolism, including PPAR signaling and bile secretion. (f) Key transcription factors, including SREBF1 and PPARA, are implicated in regulating lipid metabolic pathways in bGH-Tg mice.

Pathway enrichment analysis of the upregulated genes revealed a pronounced involvement in immune response and inflammatory processes. Specifically, pathways related to positive regulation of cytokine production, leukocyte activation, inflammatory response, platelet activation, endocytosis, extracellular matrix organization, and neutrophil degranulation were prominently affected (Fig. 1c). These findings align with previous studies indicating that GH overexpression can lead to chronic low-grade inflammation in various tissues, including the liver[11–13]. TRRUST analysis implicated several transcription factors (TFs) as potential regulators of these pathways, including TRP53, NF-kB1 (NF-κB), IRF4, STAT1, and RFXAB, (Fig. 1d) highlighting their role in orchestrating the inflammatory response in GH-overexpressing conditions[14–16].

Furthermore, we observed an upregulation of the p21-derived senescence signature in the liver of bGH-Tg mice compared to WT controls (Fig. 1b, S1a-b). Cellular senescence, characterized by irreversible growth arrest and senescence-associated secretory phenotype (SASP), has been implicated in liver aging and dysfunction[17,18]. The observed transcriptomic signature underscores the accelerated senescence and aging phenotype associated with chronic GH exposure in the liver.

In contrast, pathway enrichment analysis of the downregulated genes highlighted perturbations in metabolic pathways critical for liver function. These included mono-carboxylic acid metabolic processes, bile secretion, fatty acid metabolism, lipid homeostasis, and PPAR signaling pathways (Fig. 1e). Disruption of these pathways is consistent with previous reports linking GH excess to impaired lipid metabolism and hepato-steatosis[19]. TRRUST analysis identified key transcription factors such as SREBF1, CLOCK, and PPARA (PPAR-α) (Fig. 1f) as potential mediators of these metabolic changes, suggesting their role in regulating lipid homeostasis under conditions of GH overexpression[20–24].

### Comparative transcriptional analysis of GH overexpression and aging effects on liver transcriptome

Given the similarities in pathologies observed between GH-Tg mice and aging conditions, we sought to compare the transcriptional profiles of GH-Tg mice with those of aging mice livers. Specifically, we compared the differential gene expression profiles (DEGs) from young (10-month-old) GH-Tg mice versus WT mice with those from old (24-month-old) versus young (10-month-old) WT mice. We identified 596 genes that exhibited overlapping differential expression patterns between these two datasets (Fig. 2a). Notably, correlation analysis revealed a significantly positive correlation (r=0.30) between the DEGs of GH-Tg mice and those of aging mice, suggesting that GH overexpression induces transcriptional changes reminiscent of those seen in natural aging (Fig. 2b).

**Figure 2.**
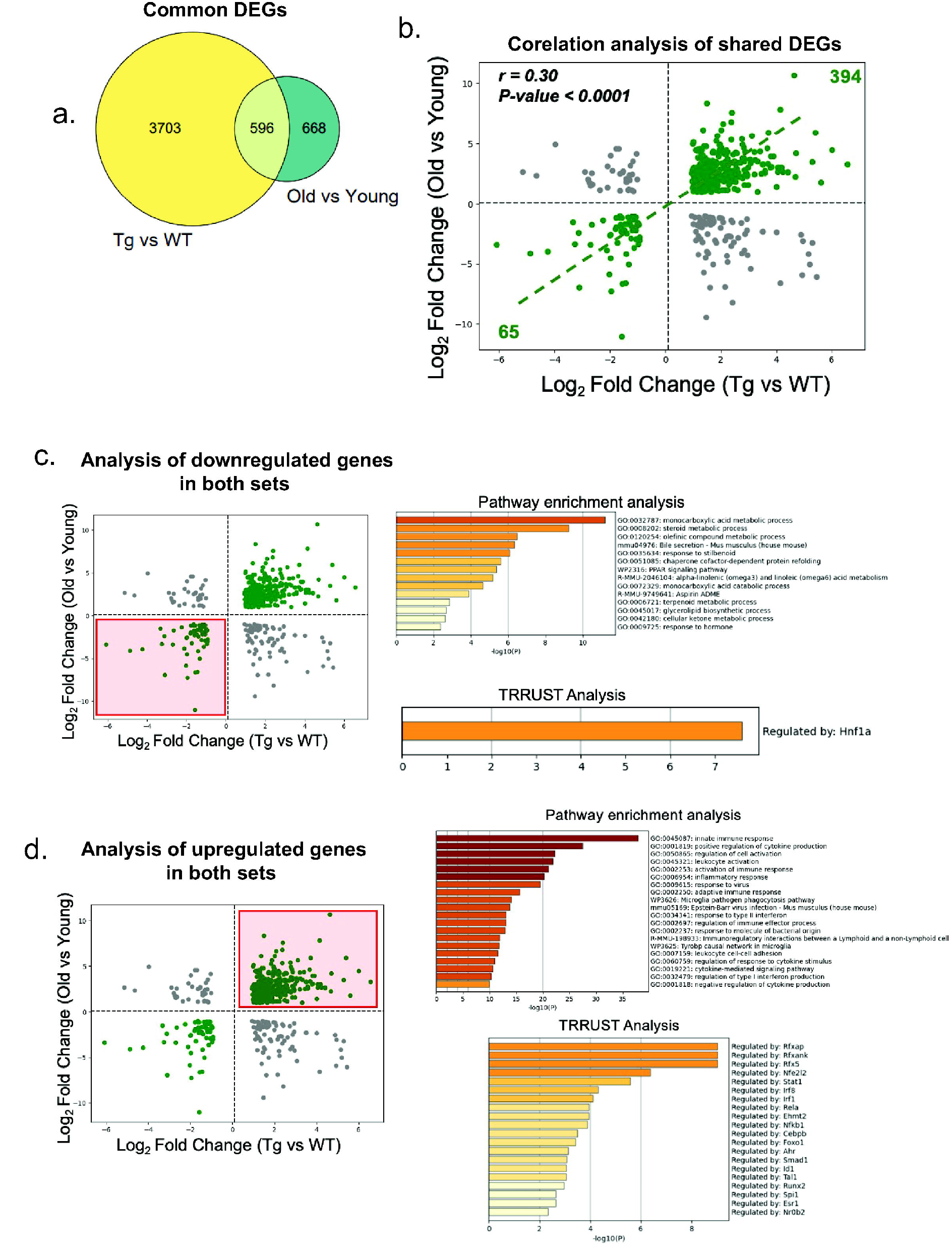
Comparative Transcriptomic Analysis of GH Overexpression and Liver Aging. (a) Venn diagram of overlapping DEGs between GH-Tg (young vs. WT) and aged liver (old vs. young WT) datasets. (b) Correlation analysis demonstrates a significant positive correlation (r=0.30) between DEGs associated with GH overexpression and liver aging. (c) Downregulated shared DEGs are enriched in metabolic pathways such as ketone metabolism and bile secretion. (d) Upregulated shared DEGs show enrichment in inflammatory pathways, including cytokine production and leukocyte activation. TRRUST analysis implicates transcription factors such as Nfe2l2 and NF-kB1 in mediating inflammatory responses common to both conditions.

Further analysis of the shared DEGs that were upregulated in both aging and GH overexpression conditions revealed enrichment in pathways associated with innate immune response, inflammatory response, leukocyte activation, and cytokine production (Fig. 2d). These findings are consistent with the concept of inflammaging, where chronic low-grade inflammation contributes to aging-related pathologies[25]. TRRUST analysis implicated transcription factors including Nfe2l2, NF-kB1 (NF-κB), RFXAP, and STAT1 (Fig. 2d) in potentially regulating these inflammatory pathways, highlighting their role in mediating the inflammatory response in both aging and GH overexpression contexts[14].

In contrast, analysis of the shared DEGs downregulated in both GH overexpression and aging conditions revealed involvement of metabolic pathways critical for liver function. These pathways included mono carboxylic acid metabolism, bile secretion, PPAR signaling pathways, and cellular ketone metabolism (Fig 2c). Disruption of these metabolic processes has been linked to impaired liver function and metabolic disorders associated with aging and GH excess[26,27]. This dual dysregulation in immune-inflammatory pathways and metabolic processes underscores the complex interplay between GH signaling and liver aging, suggesting potential therapeutic targets for mitigating age-related liver pathologies exacerbated by GH overexpression.

### Advanced glycation end products (AGEs) as downstream mediators of pathologies associated with GH overexpression

Given the observed disruption in detoxification pathways in bGH-Tg mice, we hypothesized that the pathologies associated with GH overexpression may be partially mediated by the accumulation of molecular species that induce damage over time. One such molecular insult is the accumulation of advanced glycation end products (AGEs) in various tissues[28–30]. To assess AGE accumulation, we employed mass spectrometry-based analysis of liver and serum samples from bGH-Tg mice. Our analysis revealed a significant accumulation of various types of AGEs in transgenic mice, indicating heightened glycation-related stress in these tissues (Fig. 3a-c).

**Figure 3.**
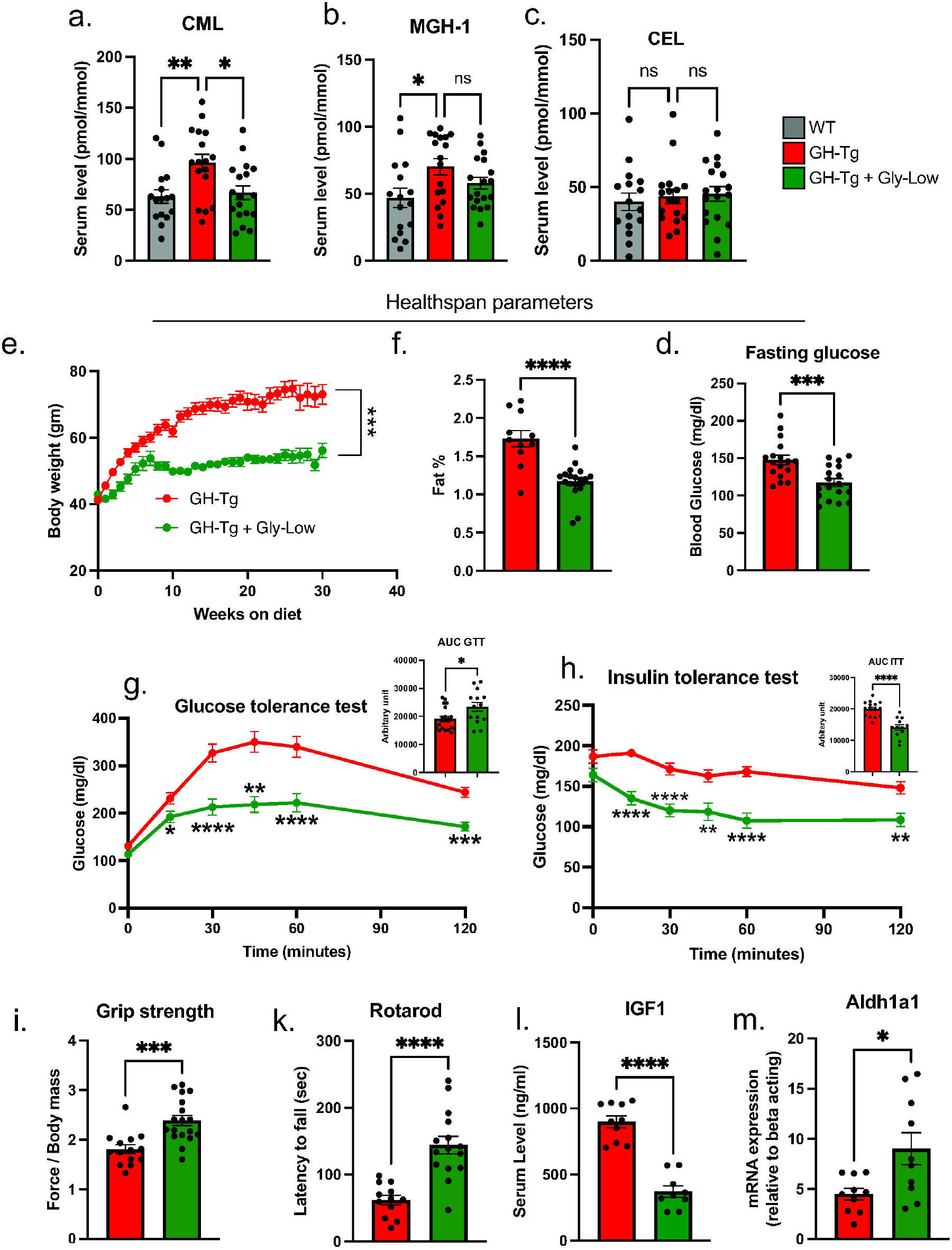
Accumulation of Advanced Glycation End Products (AGEs) and the Effect of Gly-Low Treatment in bGH-Tg Mice. (a-c) Mass spectrometry analysis shows a significant increase in AGE levels in the liver and serum of bGH-Tg mice compared to WT controls. Gly-Low treatment prevents AGE accumulation in bGH-Tg mice. (d-e) Gly-Low-treated mice exhibit reduced fat mass and improved body weight regulation compared to control-treated mice. (f-h) Improvements in fasting glucose levels, glucose tolerance, and insulin sensitivity in Gly-Low-treated bGH-Tg mice. (i-k) Functional assessments show enhanced motor coordination and grip strength in Gly-Low-treated mice. (l-m) Molecular analysis indicates reduced IGF1 expression and increased aldh1a1 expression in the livers of Gly-Low-treated mice, suggesting a reversal of glycation-induced metabolic stress. Results are represented in mean +/-SEM. Significant differences are indicated: * *p* ≤ 0.05, ** *p* ≤ 0.005, *** *p* ≤ 0.0005, **** *p* < 0.0001.

Given the pervasive role of AGEs in multiple age-related pathologies[28,31–33], we hypothesized that the accumulation of AGEs contributes to the pathologies observed in GH-Tg mice. To investigate this hypothesis, we treated GH-Tg mice with our glycation-lowering cocktail (Gly-Low), previously demonstrated to selectively target glycation-induced stress in the body[34]. As expected, long-term treatment with Gly-Low led to prevention of AGEs accumulation in the bGH-Tg mice (Fig. 3a-c). Remarkably, we observed improvement of multiple health parameters in the Gly-low treated mice compared to the control treatment, specifically, Gly-Low-treated mice exhibited rescue in body weight gain characterized by reduced fat mass percentage (Fig. 3d-e) while maintaining lean mass (data not shown). Additionally, these mice showed enhanced glucose tolerance and improved insulin sensitivity, as well as lower fasting glucose levels compared to control mice (Fig. 3d-h).

Moreover, functional assessments revealed that Gly-Low-treated mice exhibited improved motor coordination and muscle strength, as demonstrated by better performance in rotarod and grip strength assays compared to control mice (Fig. 3i-k). Interestingly, molecular analyses of liver tissues from Gly-Low-treated mice showed decreased expression levels of IGF1, a marker associated with GH overexpression-induced pathologies (Fig. 3l). Conversely, there was enhanced expression of aldh1a1 (Fig 3m), an enzyme involved in detoxification processes, suggesting a potential mechanism through which Gly-Low treatment mitigates the adverse effects of GH overexpression.

### Supplementation with Gly-Low partly reverses the transcriptomic signature in the liver induced by GH overexpression

To further elucidate the molecular effects of Gly-Low treatment on liver transcriptome in bGH-Tg mice, we hypothesized that Gly-Low would mitigate AGE-induced glycation stress and consequently reverse some of the transcriptomic alterations induced by GH overexpression. To test this hypothesis, we performed the bulk RNAseq analysis of the liver from bGH-Tg mice treated with control and Gly-Low diet. Transcriptomic analysis revealed significant segregation of different groups with 235 Differential expressed genes (Fig S2a-b). Pathway enrichment and TRRUST analysis revealed Gly-Low may regulate key liver pathways such as gluconeogenesis and cAMP-mediated signaling potentially by modulating PPARA signaling (Fig Sc-d). Next, we compared the differentially expressed genes (DEGs) from Gly-Low-treated transgenic mice (transgenic mice treated with Gly-Low vs. transgenic mice treated with a control diet) with DEGs identified in transgenic-induced changes (transgenic liver vs. WT liver). Our analysis revealed 163 shared DEGs between these datasets (Fig. 4a). Notably, these shared genes exhibited a significant negative correlation (r = -0.50), suggesting that Gly-Low treatment partially reverses the transcriptomic changes induced by GH overexpression (Fig. 4b).

**Figure 4.**
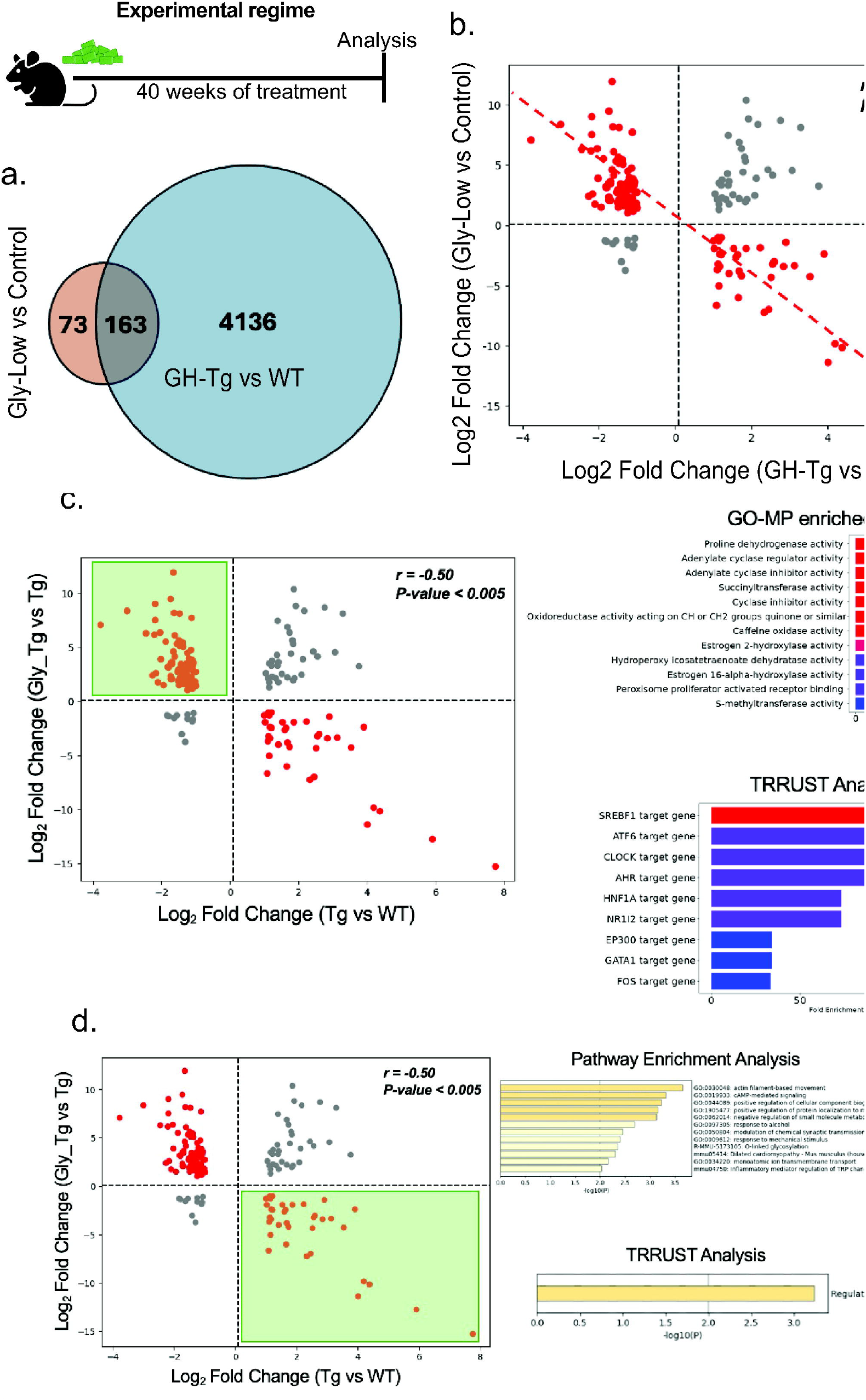
Gly-Low Treatment Partially Reverses Transcriptomic Alterations Induced by GH Overexpression (a) Venn diagram of shared DEGs between Gly-Low-treated bGH-Tg mice and GH-Tg mice (control-treated vs. WT liver). (b) Correlation analysis reveals a significant negative correlation (r=-0.50) between transcriptomic changes induced by GH overexpression and Gly-Low treatment. (c) Pathway enrichment analysis of genes upregulated by Gly-Low treatment identifies processes such as adenylate cyclase activity and detoxification pathways. (d) Downregulated pathways include inflammatory mediator regulation and actin filament-based movement. TRRUST analysis implicates transcription factors such as PPARA and CLOCK as regulators of these transcriptomic reversals.

In-depth pathway enrichment analysis of the genes upregulated in GH-Tg liver, but downregulated following Gly-Low treatment, identified key pathways such as actin filament-based movement, cAMP-mediated signaling, response to alcohol, O-linked glycosylation, and inflammatory mediator regulation. TRRUST analysis implicated PPARA as a potential regulator of these pathways (Fig. 4d). The role of PPARA in lipid metabolism and inflammation is well-documented, suggesting that Gly-Low’s beneficial effects may be mediated through modulation of PPARA activity[23,35].

Conversely, genes that were downregulated in GH-Tg mice and upregulated after Gly-Low treatment were associated with pathways involving proline dehydrogenase activity, adenylate cyclase activity, and estrogen 2-hydroxylase activity (Fig. 4c). TRRUST analysis identified transcription factors such as SREBF1, ATF6, and CLOCK as key regulators of these processes (Fig. 4c). SREBF1 and ATF6 are critical for lipid homeostasis and endoplasmic reticulum stress response, respectively [36,37]. CLOCK, a core circadian regulator, has been shown to influence detoxification pathways and metabolic processes, indicating that modulation of circadian genes might be a viable strategy to combat glycation stress[38–40].

In summary, these findings indicate that Gly-Low partly counteracts GH-induced transcriptomic alterations, predominantly by reversing changes in key metabolic and inflammatory pathways. This underscores the therapeutic potential of targeting AGE-induced glycation stress and circadian gene modulation to mitigate pathologies associated with chronic GH overexpression.

## Discussion

This study provides significant insights into the impact of chronic growth hormone (GH) overexpression on liver metabolism, highlighting key changes that may drive GH-induced metabolic disorders. Using a bovine GH-overexpressing transgenic (bGH-Tg) mouse model, we observed marked transcriptomic changes, highlighting dysregulation in fatty acid metabolism, heightened inflammatory responses, and evidence of cellular senescence within the liver, resembling aging profiles. These findings support the hypothesis that GH excess not only affects liver metabolism directly, but also accelerates hepatic aging processes, contributing to systemic metabolic dysregulation. The suggested acceleration of hepatic aging is consistent with the evidence that GH transgenic mice have markedly reduced longevity along with phenotypic traits that have been interpreted as symptoms of early aging[41,42]. Interestingly, mice, in which GH signaling is suppressed rather than enhanced, are characterized by extended longevity and slower and/or delayed aging.[43–46]

Our transcriptomic analysis revealed that GH overexpression perturbs immune and inflammatory pathways, corroborating previous findings linking GH excess to chronic low-grade inflammation. The enriched inflammatory pathways, including leukocyte activation and cytokine production, suggest that GH-driven inflammaging may be mediated by key transcription factors such as NF-kB and STAT1. This pattern of inflammatory upregulation parallels liver aging, suggesting that GH overexpression may act as an aging accelerant through sustained inflammation. Such a mechanism aligns with the concept of inflammaging, where chronic, low-grade inflammation contributes to metabolic and functional decline in aging tissues.

In parallel, GH overexpression disrupts metabolic pathways central to liver function, including fatty acid oxidation, bile secretion, and lipid homeostasis. These disruptions mirror changes observed in aged livers and underscore the liver’s sensitivity to GH dysregulation. Transcription factors such as PPAR-α, SREBF1, and CLOCK were implicated as central regulators of these pathways, highlighting their role in maintaining metabolic stability under normal conditions and their potential dysregulation in GH-induced hepatic stress. The overlap in transcriptional changes between GH-Tg mice and aged livers further reinforces the idea that GH overexpression accelerates hepatic aging, likely through combined inflammatory and metabolic stress.

Another novel aspect of this study is the identification of advanced glycation end products (AGEs) as a prominent feature in the livers of bGH-Tg mice. The accumulation of AGEs, compounds linked to aging and metabolic diseases, suggests that GH-induced stress may overwhelm hepatic detoxification pathways, promoting glycation stress. AGEs are known to contribute to cellular dysfunction by altering protein structure and triggering inflammatory signaling, potentially exacerbating GH-driven liver pathologies. These findings position AGEs as not only a marker of GH-induced metabolic stress but also a possible mediator of GH-related liver dysfunction.

Our findings indicate that glycation-lowering compounds (Gly-Low) can mitigate many of the deleterious effects associated with GH overexpression. Gly-Low treatment reversed several GH-induced transcriptomic changes, particularly in pathways related to inflammation and metabolic homeostasis. Notably, Gly-Low downregulated inflammatory pathways and upregulated detoxification and lipid metabolic pathways, suggesting that glycation reduction can restore, at least partially, liver-mediated functions and glucose homeostasis disrupted by GH excess. This therapeutic approach has potential implications for managing conditions associated with chronic GH dysregulation, such as acromegaly, and may offer broader benefits for liver pathologies where glycation stress is prevalent.

In conclusion, this study underscores the profound impact of GH overexpression on liver aging and metabolic disorders, with AGEs identified as potential targets for therapeutic intervention. Glycation-lowering strategies may serve as effective treatments for alleviating GH-induced metabolic and inflammatory disruptions in the liver, offering a promising avenue for addressing age-related metabolic diseases associated with GH dysregulation.

## Methods

### Ethics Statement

The guidelines of the Southern Illinois University Institutional Animal Care and Use Committee were followed in the performance of this work.

### Animals

For this research, bovine GH transgenic mice were created in our colony from animals kindly donated by Dr. T. Wagner and J. S. Yun of Ohio University in Athens, Ohio[47]. Male transgenic mice were mated with normal (C57BL6/J x C3H/J F1) hybrid females to produce the mice. For this experiment, normal siblings of bGH Tg mice served as controls. Animals were kept in climate-and light-controlled environments (21-23°C with a 12-hour cycle of light and darkness). The Institutional Animal Care and Use Committee at Southern Illinois University approved all animal methods used in this investigation. The total number of mice used in this study was around 150, i.e., 15-20 mice per group.

### Weaning and genotype confirmations

All animals were weaned 21-22 days after birth and housed in a cage of 4-5 animals per group. Though PEPCK-bGH mice are phenotypically distinct, being large from their normal littermates, tissue genotyping was done via normal PCR and qPCR, and the presence of the bovine growth hormone gene was confirmed. The PCR products of the bGH gene, with a size of 446 bp, were amplified using bGH primers. The DNA amplicons were detected in 1.5% agarose gel following electrophoresis for about 1 hour and visualized under UV light.

### Body weight measurements

Measurement of body weight was done once a week, starting at the 10th week to the 40th week.

### Normal diet and CRM diet composition

Mice were fed either a standard low-fat chow diet (21% fat (kcal), 60% carbohydrate (kcal) Envigo: TD.200743), a standard high fat chow diet (60% fat (kcal), 21% carbohydrate (kcal), Envigo: TD.200299), a standard low-fat chow diet supplemented with our Gly-Low compound cocktail (21% fat (kcal), 60% carbohydrate (kcal) Envigo: TD.200742), or a standard high fat chow diet supplemented with our Gly-Low compound cocktail (60% fat (kcal), 21% carbohydrate (kcal).

A combination of supplemental grade compounds, safe to be consumed in set dosages, were prepared and incorporated into a modified pre-irradiated standard AIN-93G mouse chow diet from Envigo. The cocktail consists of alpha lipoic acid (20.19%), nicotinamide (57.68%), thiamine hydrochloride (4.04%), piperine (1.73%), and pyridoxamine dihydrochloride (16.36%), and is supplemented in the diet to achieve a daily consumption rate in mg/kg of body weight/day. For diet, these percentages translate to 3 g/kg alpha lipoic acid, 8.57 g/kg nicotinamide, 0.6 g/kg thiamine hydrochloride, 0.26 g/kg piperine, and 2.43 g/kg pyridoxamine dihydrochloride.

### Insulin tolerance test

Insulin sensitivity test was done at two time points, 19th and 37th weeks of mice age, i.e., 9 and 27 weeks after starting the CRM treatment via tail bleed in non-fasted mice. Insulin (Sigma, St. Louis, MO, USA) was injected intraperitoneally at one international unit (IU) per kg body weight. Blood glucose was measured at 0, 15, 30, 45, 60, and 120 minutes via glucometer (Agal Matrix). Data from the ITT are shown as a percentage of baseline glucose and as an area under the curve.

### Glucose tolerance test

One week after ITT, a glucose tolerance test was done at two-time points, the 20th and 38th weeks of mice age, i.e., 10 and 28 weeks after starting the CRM treatment via tail bleed. All mice were fasted overnight for 16 hours before the glucose injection. Glucose was injected via intraperitoneal injection at 2 grams/kg body weight. Blood glucose was measured at 0, 15, 30, 45, 60, and 120 minutes via glucometer (Agal Matrix). Data from the GTT are shown as a percentage of baseline glucose and as an area under the curve.

### Rotarod

A rotarod test was done to assess the neuromuscular balance or motor coordination at week 39. Mice were trained to run on a rotating wheel for 5 minutes two times on day 1, and an actual run was done on day 2 to calculate the average latency for falling.

### Grip-Strength

Grip strength test was done on 39-40th weeks to determine the muscular strength generated by forearm muscle in 3 trials on the same day. The body weight normalized all grip strength values.

### Tissue Collection

All mice were euthanized between 9 AM and 11 AM during the 40 weeks of age or 30th week after treatment to collect the blood and tissues, including liver, muscle, pancreas, colon, heart, brain, hypothalamus, hippocampus, subcutaneous, epididymal, brown and peri-renal fat. Isoflurane was used to anesthetize the animals before being bled by heart puncture and decapacitated. During the collection, tissues were frozen in dry ice and stored at -80°C.

### RNA Extraction and cDNA Synthesis

Total RNA was extracted utilizing the Trizol method, and cDNA was produced via reverse transcription. Two-step real-time PCR system was used to perform SYBR green-based real-time reverse transcriptase PCR. The housekeeping gene was selected based on the type of tissue used to normalize values for the genes of interest, and the relative amounts of mRNA were calculated using the Ct technique.

### RNA sequencing

RNA was isolated using Zymo research quick RNA miniprep kit (cat # 11-328) according to the manufacturer’s recommendations. Isolated RNA was sent for library preparation and sequencing by Novogene Corporation Inc. where RNA was poly-A selected using poly-T oligo-attached magnetic beads, fragmented, reverse transcribed using random hexamer primers followed by second strand cDNA synthesis using dTTP for non-directional library preparation. Samples underwent end repair, A-tailing, adapter ligated, size selected, amplified, and purified. Illumina libraries were quantified using Qubit and qPCR and analyzed for size distribution using a bioanalyzer. Libraries were pooled and sequenced on an Illumina Novoseq 6000 to acquire paired-end 150 bp reads. Data quality was assessed, and adaptor reads and low-quality reads were removed. Reads that passed the quality filtering process were mapped paired end to the reference genome (GRCm38) using Hisat2 v2.0.5. feature Counts v1.5.0-p3 was used to count reads that mapped to each gene. Differential expression analysis was performed using DESEq2 (1.20.0). Where indicated, bootstrapping was performed using R (R version 4.1.2) program ‘boot’ (1.3-28.1). To determine the expected mean and standard deviation, n=i log2 fold changes were randomly selected 1000 times, in which i is the number of genes in the gene set.

### Protein analysis

The blood sample was taken during the sacrifice and centrifuged to get plasma. According to the manufacturer’s instructions, ELISA was used to quantify the serum levels of IGF-1, and insulin (Crystal Chem Inc).

### Quantification of Di-carbonyls

200 μL of 80:20 MeOH:ddH_2_O (−80°C) containing 50 pmol ^13^C_3_-MGO was added to 10 μL of serum and extracted at −80°C overnight. Insoluble protein was removed via centrifugation at 14,000 x g for 10 min at 4°C. Supernatants were derivatized with 10 μL of 10 mM o-phenylenediamine for 2 h with end-over-end rotation protected from light. Derivatized samples were centrifuged at 14,000 x *g* for 10 min, and the supernatant was chromatographed using a Shimadzu LC system equipped with a 150 × 2mm, 3μm particle diameter Luna C_18_ column (Phenomenex, Torrance, CA) at a flow rate of 0.450 mL/min. Buffer A (0.1% formic acid in H_2_O) was held at 90% for 0.25 min, then a linear gradient to 98% solvent B (0.1% formic acid in acetonitrile) was applied over 4 min. The column was held at 98% B for 1.5 min, washed at 90% A for 0.5 min, and equilibrated to 99% A for 2 min. Multiple reaction monitoring (MRM) was conducted in positive ion mode using an AB SCIEX 4500 QTRAP with the following transitions: *m/z* 145.1→77.1 (MGO); *m/z* 235.0→157.0 (3-DG); m/z 131.0→77.0 (GO); m/z 161.0→77.0 (HPA); m/z 148.1→77.1 (^13^C_3_-MGO, internal standard).

### Statistical Analysis

The data are shown as mean ± SEM. GraphPad Prism Software V8.0 was used for the analysis. A student’s t-test was used to compare the means of the two groups. The significance of the differences in the insulin tolerance test (ITT) and glucose tolerance test (GTT), indirect calorimetry was calculated by comparing the area under the curve (AUC). Two-way or three-way ANOVA with Tukey’s post hoc tests were used to compare the mean for more than two groups. P values less than 0.05 were considered statistically significant. Significant differences are indicated: * *p* ≤ 0.05, ** *p* ≤ 0.005, *** *p* ≤ 0.0005, **** *p* < 0.0001.

## Author contributions

PS, AG, LH, JG, AB, PK were involved in conceptualization and supervision. PS, LH, PK, AB drafted the manuscript. PS, AG, LH were involved in analysis and visualization. PS, AG, LH, MT were involved in experimental procedures and live animal experimentation. All authors have read and agreed to the final version of the manuscript.

## Funding statement

We acknowledge support from the National Institute of Health: R01AG038688 (PK), R01AG068288 (PK), R01AG061165 (PK), the Larry L. Hillblom Foundation (PS), the Hevolution Foundation HF-GRO-23-1199066-37 (AB), the William E. McElroy Charitable Foundation (AB), and the SIU Geriatrics Initiative (AB).

## Conflict of interest statement

Authors have no conflicts of interest to declare.

## Data availability

The data supporting this study’s findings are available from the corresponding author upon request.

## Notes

### Competing Interest Statement

The authors have declared no competing interest.

