## Supplementary material for "Growth Hormone Excess Drives Liver Aging via increased Glycation stress": Suppli Figure S1

### a. MCODE analysis of upregulaged DEGs

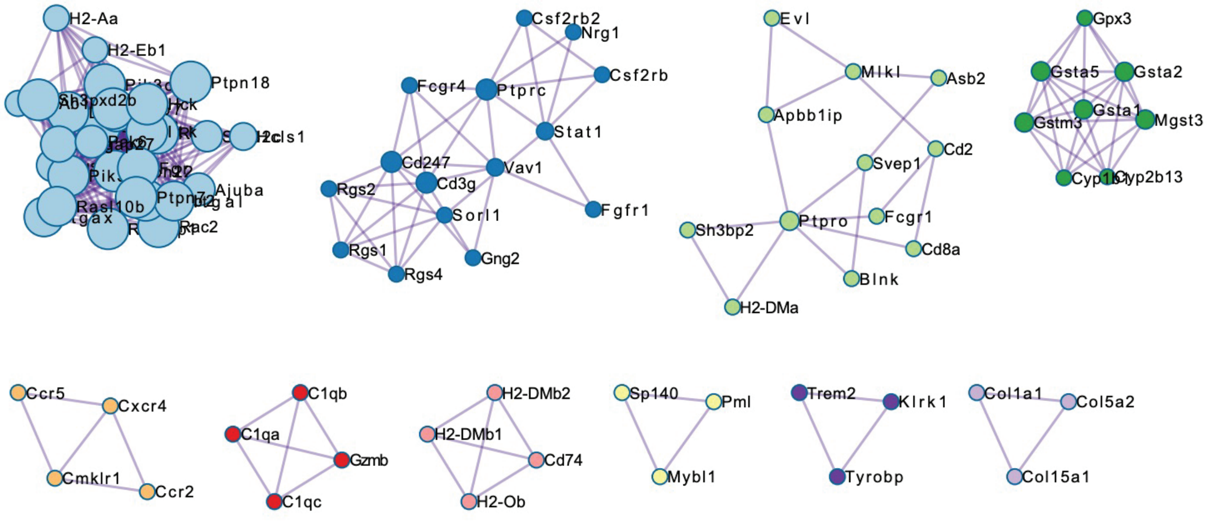


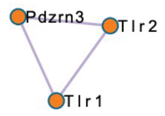


| Color | MCODE | GO | Description | Log10(P) |
|---|---|---|---|---|
| | MCODE_1 | WP85 | Focal adhesion | -13.5 |
| | MCODE_1 | GO:1903039 | positive regulation of leukocyte cell-cell adhesion | -9.8 |
| | MCODE_1 | GO:0022409 | positive regulation of cell-cell adhesion | -9.2 |
| | MCODE_2 | R-MMU-2029482 | Regulation of actin dynamics for phagocytic cup formation | -7.3 |
| | MCODE_2 | R-MMU-202427 | Phosphorylation of CD3 and TCR zeta chains | -7.2 |
| | MCODE_2 | R-MMU-2029481 | FCGR activation | -7.1 |
| | MCODE_3 | mmu04640 | Hematopoietic cell lineage - Mus musculus (house mouse) | -6.8 |
| | MCODE_3 | GO:0002252 | immune effector process | -5.1 |
| | MCODE_3 | GO:0046649 | lymphocyte activation | -4.9 |
| | MCODE_4 | mmu00480 | Glutathione metabolism - Mus musculus (house mouse) | -13.5 |
| | MCODE_4 | mmu00980 | Metabolism of xenobiotics by cytochrome P450 - Mus musculus (house mouse) | -13.4 |
| | MCODE_4 | mmu05204 | Chemical carcinogenesis - DNA adducts - Mus musculus (house mouse) | -13.0 |
| | MCODE_5 | GO:0019886 | antigen processing and presentation of exogenous peptide antigen via MHC class II | -12.1 |
| | MCODE_5 | GO:0002495 | antigen processing and presentation of peptide antigen via MHC class II | -11.9 |
| | MCODE_5 | GO:0002504 | antigen processing and presentation of peptide or polysaccharide antigen via MHC class II | -11.8 |
| | MCODE_6 | R-MMU-173623 | Classical antibody-mediated complement activation | -10.1 |
| | MCODE_6 | GO:0098883 | synapse pruning | -9.8 |
| | MCODE_6 | GO:0150146 | cell junction disassembly | -9.4 |
| | MCODE_7 | GO:0007204 | positive regulation of cytosolic calcium ion concentration | -8.1 |
| | MCODE_7 | WP189 | GPCRs class A rhodopsin like | -7.9 |
| | MCODE_7 | GO:0006935 | chemotaxis | -7.2 |
| | MCODE_9 | R-MMU-8948216 | Collagen chain trimerization | -8.4 |
| | MCODE_9 | R-MMU-2022090 | Assembly of collagen fibrils and other multimeric structures | -8.0 |
| | MCODE_9 | R-MMU-1442490 | Collagen degradation | -7.8 |
| | MCODE_10 | R-MMU-2424491 | DAP12 signaling | -8.6 |
| | MCODE_10 | R-MMU-2172127 | DAP12 interactions | -7.6 |
| | MCODE_10 | GO:0002705 | positive regulation of leukocyte mediated immunity | -6.1 |

### b. Gene set enrichment analysis with Tabula Muris senescence database

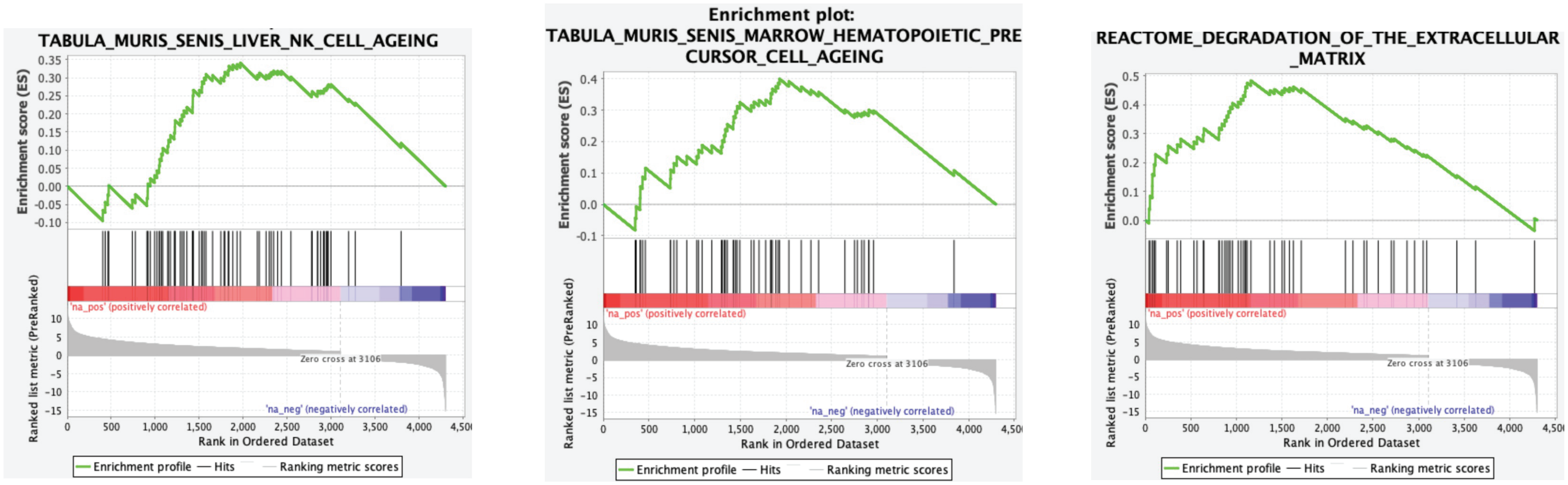


Fig S1
