## Supplementary figures and images for "Growth Hormone Excess Drives Liver Aging via increased Glycation stress"

### Suppli Figure S2

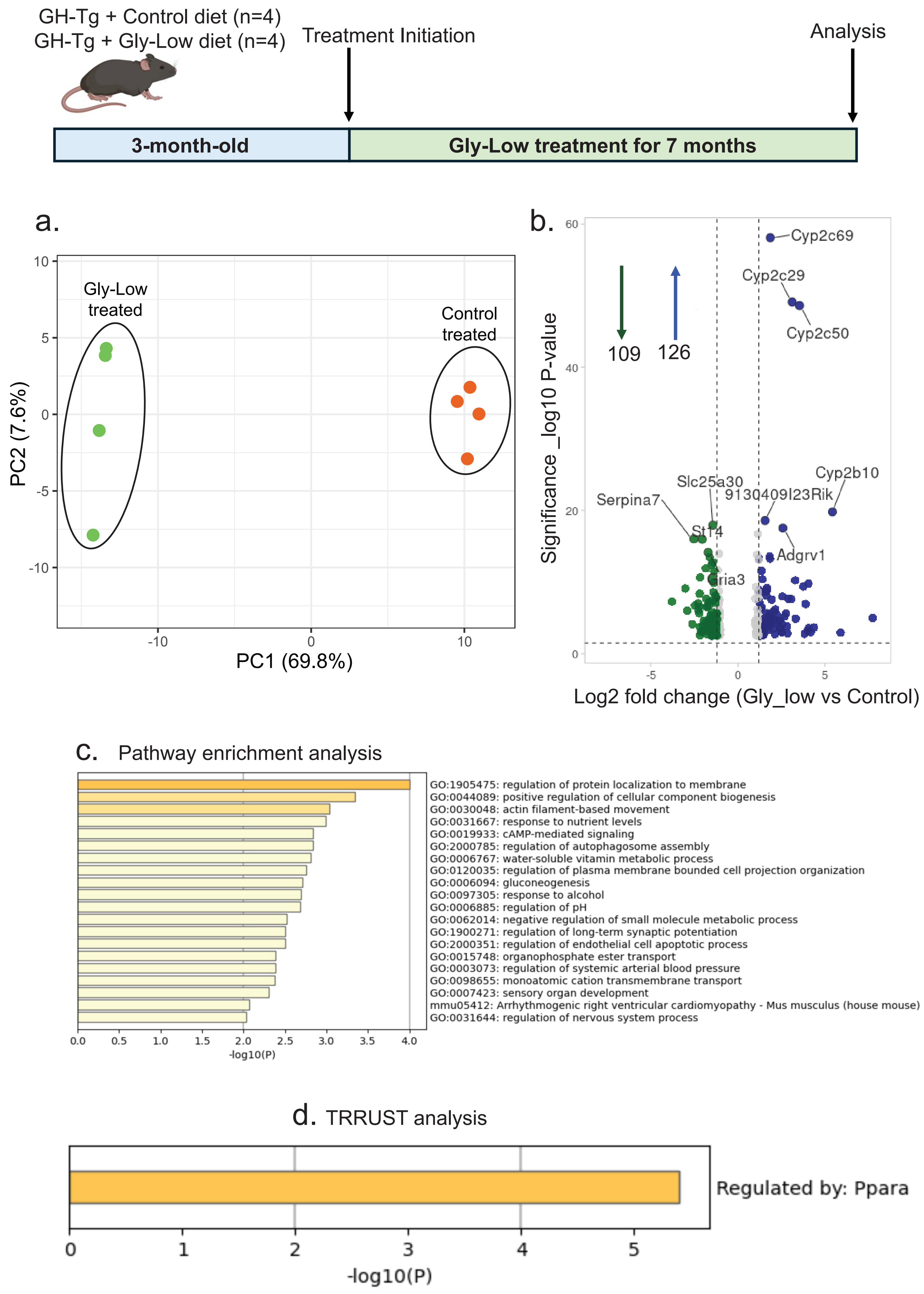
